# Spike synchrony as a measure of Gestalt structure

**DOI:** 10.1101/2023.12.05.570259

**Authors:** Viktoria Zemliak, Julius Mayer, Pascal Nieters, Gordon Pipa

## Abstract

The function of spike synchrony has long been debated in the neuroscience community. Some researchers view synchrony as a byproduct of brain activity, while others argue that it serves as a mechanism for binding perceptual features. We, however, argue for an alternative view on the computational role of synchrony: it can serve as a mechanism for estimating the prior probability of incoming stimuli. In V1, this probability can be estimated by comparing the input stimulus with previously acquired visual experience. Such experience is encoded in horizontal intracortical connections, as they are subject to change, learn, and adapt to incoming visual stimuli throughout the course of life. Thus, the V1 connectivity structure can encode the acquired visual experience in the form of its aggregate statistics. Since the aggregate statistics of natural images tend to follow the Gestalt principles, we can assume that V1 is more often exposed to Gestalt-like stimuli, and this is manifested in its connectivity structure. At the same time, the connectivity structure has an impact on the synchrony of neuronal firings in V1. In our work, we used a spiking model with V1-like connectivity to demonstrate that spike synchrony reflects the Gestalt structure of the stimulus. We conducted simulation experiments with three Gestalt laws: proximity, similarity, and continuity, and found significant differences in firing synchrony for stimuli with varying degrees of Gestalt-likeness. This allows us to conclude that spike synchrony indeed reflects the Gestalt structure of the stimulus, which can be interpreted as a mechanism for prior probability estimation.

**Author Summary:** Neurons in the brain fire synchronously in different contexts, in response to different stimuli. However, the function of synchrony is very debatable in the neuroscience community. We suggest that synchrony plays an important role in brain computations: it can estimate how well a network is familiar with the incoming stimulus. We tested our hypothesis on the simulation of neuronal activity in a model of primary visual cortex - a brain region which is processing simple visual features, such as lines of different angle orientation. The connections between neurons in this brain area store the acquired visual experience in the form of its statistics, such as edge cooccurrences. We built the connectivity in our model in accordance with principles of connectivity in primary visual cortex, assuming that it reflects natural images statistics. We tested our model on artificial stimuli, which represented more and less experienced visual features. Our results showed that spike synchrony is a global phenomenon which emerges across the large parts of the network, and it reflects the familiarity of the network with the stimulus. Thus, it can serve as a form of recognition memory and has a potential for usage in various applications which require estimation of input familiarity.

## Introduction

Neurons in various cortical areas fire synchronously in response to various tasks. First evidence of synchronous firing was found in the sensory cortex, e.g. the primary visual cortex [1], extrastriate [2] and primary auditory cortex [3, 4]. Additionally, spike synchrony was later discovered in the primary motor cortex [5] and frontal cortex [6].

However, the function of spike synchrony has been a subject of a long-standing debate in the neuroscience community. Two most prominent hypotheses argue that synchrony is either a byproduct of brain activity [7, 8], or that it serves as a mechanism of binding the perceived information, thus creating a representation of the whole from the separate input components [9, 10]. Along with the binding hypothesis, some research suggests that synchrony in the beta or gamma frequency range can serve for attention modulation and relevant stimulus selection [11–13]. Synchrony associated with the attention distribution can also be observed between multiple functional brain areas [14].

In our work, we argue for an alternative view on the computational role of synchrony: it can indicate the Gestalt structure and hence the prior probability of the incoming visual stimulus [15]. That is, if the cortical units are familiar with the stimulus, they tend to fire synchronously in response to it. We suggest that spike synchrony in populations of neurons is an estimator of the match between the stimulus and the stimulus prior that is encoded in the network structure.

Additionally, we argue that synchrony, and in turn the estimate of the stimulus matching and the encoded prior, is an emergent property that spreads across larger parts of the network. Spike synchrony provides additional information which is not directly encoded in the stimulus pattern. Last but not least, we want to highlight that we do not rely on oscillatory activity patterns and their frequency or phase. Instead, we show that the computational role of synchrony arises solely from the noise induced coherence - a mechanism depending on the network connectivity structure.

### Synchrony evidence

Spike synchrony has been observed in various behavioral contexts. In the sensory cortex, synchronous firing depends on properties of the input. For example, neighboring neurons of the primary auditory cortex with similar receptive fields fire simultaneously, which can be seen at all layers between all distinguishable cell types [16]. The synchronous firing of the distant neurons was shown in the primary somatosensory cortex [17]. Spike synchrony has been widely observed in the primary visual cortex (V1). It depends on certain geometrical characteristics of the input stimuli: spatial continuity [1] and orientation similarity [18]. Thus, neurons of V1 fire synchronously in response to continuous visual stimuli and stimuli with homogeneous angle orientation.

Besides the sensory cortex, synchronous firing was found in the executive areas. In the primary motor cortex, cells with similar muscle fields tend to fire synchronously [19]. There was also evidence that neurons of the primary motor cortex fire synchronously with both the performed and the intended action [20]. The results of the latter might be perceived controversial due to the choice of the analysis method, but they were replicated with use of the more robust techniques [21].

Synchronous firing can also correlate to higher cognitive functions. Cells in the middle temporal area (MT) fire synchronously in relation to decision making [22]. In the prefrontal cortex, the neurons can exhibit simultaneous firing during working memory tasks [23, 24]. The firing of the neurons of the inferotemporal cortex can synchronize when monkeys process face features [25].

### Neural basis of synchrony

The neural basis of spike synchrony lies in horizontal intracortical connections [26–28]. These connections mostly arise from pyramidal cells and are parallel to the cortical surface [29]. Their important property is that neurons which fire in similar contexts tend to form such strong connections. Horizontal connections were found in various brain areas, e.g. in the primary motor cortex between neurons representing similar muscle groups [30].

The sensory cortex was also shown to be full of intracortical horizontal connections. For example, cells of the auditory cortex that have similar bandwidth and characteristic frequency selectivity are well connected to each other [31]. Horizontal connections in the somatosensory cortex can be formed even between the distant cells which respond to tactile stimuli at the opposing fingertips [32]. Interestingly, in this case well connected cells of the somatosensory cortex do not have similar receptive fields, but rather are engaged in the common somatosensory context.

Since the current research is primarily focused on the primary visual cortex, we describe its horizontal connections and other architectural properties in more detail. V1 area is known to be retinotopically organized, i.e. its neurons respond to the stimuli, whose retinal coordinates match the cortical position of these neurons. Besides, these neurons have orientation selectivity. Thus, each V1 cell is spatially and orientationally tuned: it fires when there is a visual stimulus with the particular angle in the particular region of the retina. The cells that are responding to the specific orientation are forming cortical columns, which are in turn organized in hypercolumns. Columns that constitute one hypercolumn share the similar receptive fields [33].

There are two main factors that influence presence and strength of the horizontal intracortical connections in V1: the spatial proximity of the receptive fields, and the similarity of the preferred orientation [34, 28]. Thus, the cells that are responding to spatially neighboring visual stimuli of the similar orientation, tend to have strong horizontal connections. On the physiological level, the stronger connections between neurons with similar orientation tuning are manifested in larger synapses [35].

At the same time, intracortical horizontal connections are subject to change, learn, and adapt to the experience. In adults, the plasticity of horizontal intracortical connections is associated with acquiring new skills [36]. During the development of the visual cortex, first the unclustered and weakly specified horizontal connections arise. Later they become more refined and fine-tuned under the control of visual experience [37, 38].

### Natural image statistics and Gestalt principles

Thus, the horizontal connectivity in V1 reflects the acquired visual experience. This experience can be manifested in the form of aggregate statistics of experienced natural visual stimuli [39]. There has been numerous research on various environmental statistics that can influence neuronal connectivity and neuronal activity patterns (see [40] for review). So, what are the relevant statistics of natural visual scenes that shape the horizontal connectivity in V1?

Brunswik & Kamiya 1953 have first shown that one of the crucial statistical bases in natural images are the Gestalt principles [41]. The Gestalt principles describe the mechanism for grouping and interpreting visual elements. The elements which are following these principles are more likely to be perceived as a holistic object, rather than separate ones. The Wertheimer landmark paper included six Gestalt principles: proximity, similarity, uniform density, direction, common fate, good continuation [42]. Since then, the Gestalt theory has been highly debatable, and various researchers have suggested up to a hundred new grouping principles [43]. On the contrary, there is an idea that all Gestalt principles are the special instances of a single general Good Gestalt principle [44].

The natural images statistics have been shown to be consistent with certain Gestalt principles, including proximity, continuity and similarity [41, 45–48]. It was also demonstrated how the Gestalt-based perceptual grouping can be shaped and modulated by the visual experience [49, 50]. Sigman et al., 2001 found that the regularities in natural images follow the Gestalt continuity principle, and showed how such regularities are reflected in the V1 connectivity [48]. In Onat, Jancke, & König, 2013, the Gestalt principles were also shown to be manifested in the V1 connectivity, but with the natural movies used as stimuli, rather than static images [39].

As summarized in Korndörfer et al., 2017, most often experienced and behaviorally relevant natural visual stimuli appear to have the Gestalt-like structure, and it manifests at the level of intracortical connectivity [15]. And this Gestalt-structured connectivity, in turn, is assumed to be the basis of spike synchrony in visual cortex [26–29].

## Results

We used a pulse-coupled fully-excitatory Izhikevich spiking network [51] in a noisy environment, to show that neuronal synchrony reflects how closely the visual input stimulus follows the Gestalt principles. Our network consists of Izhikevich neurons with additional dynamics of open receptors and neurotransmitter conductances (see Methods for a detailed model description). Neurons in the model are organized retinotopically, have spatial and orientation selectivity, and are connected in all-to-all fashion. The schematic model representation is depicted in Fig 1.

**Figure 1.**
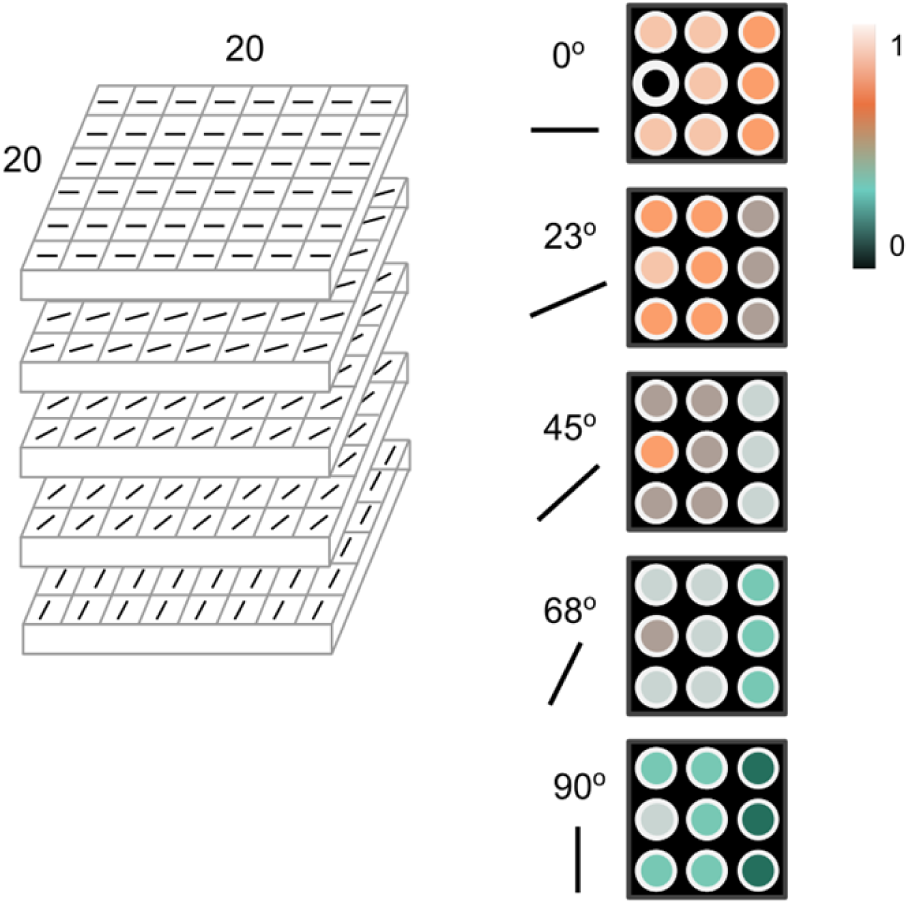
Model architecture and connectivity. **A.** The model consists of neurons with spatial and angle preferences. **B.** The example of connectivity between neurons. All connections of a black neuron with neurons in a neighborhood of 2 are shown. A color bar shows the relative connectivity strength.

The connectivity structure is defined by a connectivity matrix and follows a simple rule: the connection strength between two neurons depends on their spatial distance and difference in orientation selectivity. Thus, neighboring neurons with shared orientation preferences are strongly connected, whereas the most distant neurons with different orientation selectivity form the weakest connections. The preferential connections and larger synapses between neurons with similar receptive fields were shown in V1 [28, 35].

Thus, each neuron receives lateral input through the horizontal connections defined in a connectivity matrix, and external input from the input images. We simulated 2 seconds of activity of each neuron. Following Korndörfer et al., 2017, we measured the synchrony of their firing responses as the noise-induced coherence, and refer to it as Rsync (see Methods) [15].

To study how spike synchrony reflects the Gestalt structure of stimuli, we ran three experiments for three Gestalt principles: proximity, similarity, and continuity. Within each experiment, we first generated a set of artificial input stimuli, then ran 100 simulation trials of our model on each stimulus (14 stimuli for all experiments in total x 100 trials). One run of every trial simulated 1500 ms of neuronal activity. Finally, for each input stimulus, we measured and averaged firing synchrony over all 100 trials. Fig 2 shows the full experimental pipeline.

**Figure 2.**
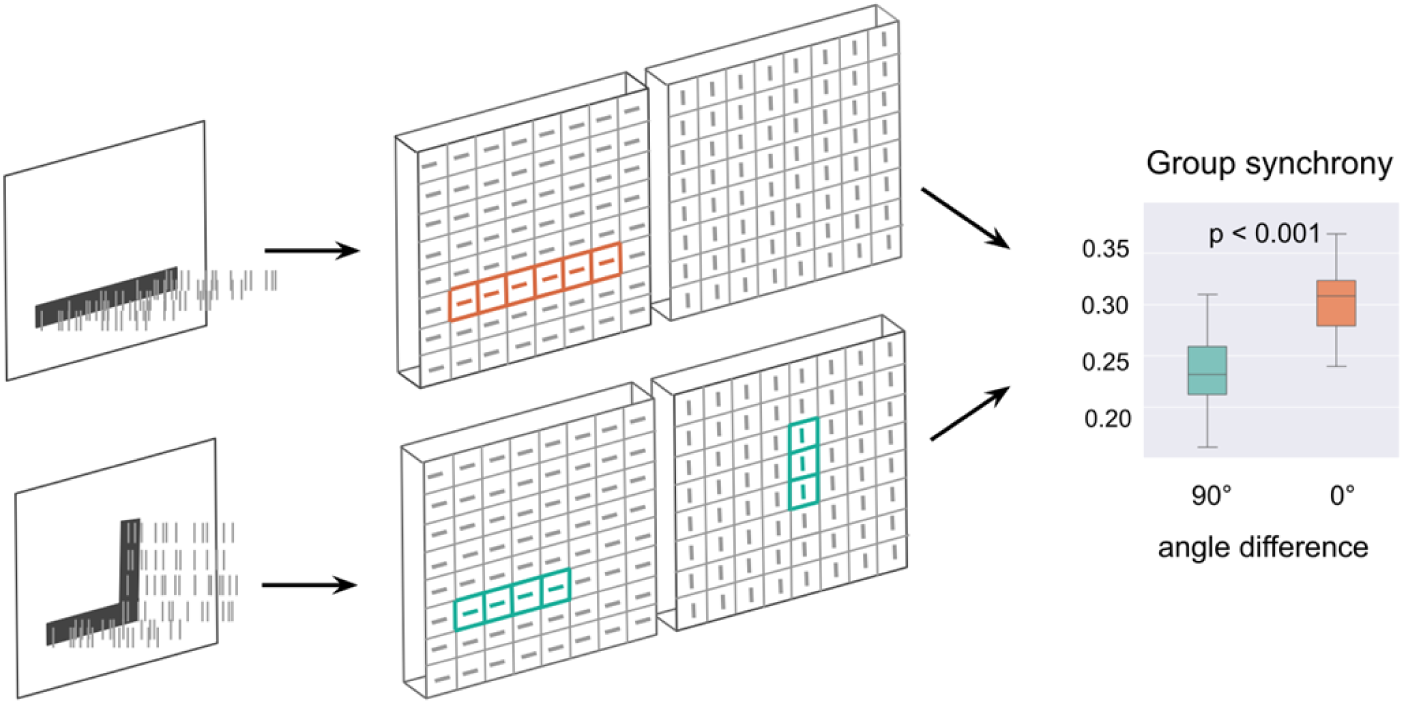
Simulation pipeline. **A.** Stimuli patterns of various Gestalt-likeness are sent to the model in the form of Poisson spike trains. **B.** Spatially- and orientation-selective neurons in the model detect angles and become activated in the locations which receive input. **C.** Spike synchrony is measured across the neurons from the activated subsets.

For each simulation experiment, we used a set of stimuli that followed the Gestalt principles to a different degree. In other words, stimuli had gradual Gestalt-likeness. One stimulus consisted of several line segments, which were organized in greater or lower accordance with a certain Gestalt principle.

1. The Gestalt principle of *proximity* implies that single visual items near each other are perceived as an aggregation into a larger visual composition. In our context, that refers to input stimuli consisting of two segments on various distances from each other (see Fig 9A in Methods). The smaller the distance between the segments, the more Gestalt-like the entire stimulus.
2. Visual components can also be integrated into perceptual groups based on the similarity of their appearance. This classification of elements by visual features is an instance of the Gestalt principle of *similarity*. In our work, we interpret similarity as shared angle orientation: if two parts of the stimulus share the similar orientation, the entire stimulus is highly Gestalt-like. Hence, the bigger the angle difference, the less Gestalt-like the stimulus is (see Fig 9B in Methods).
3. The Gestalt principle of *continuity* is based on the idea that visual elements are preferably grouped if they are organized in a continuous line. Thus, elements with abrupt changes in orientation are more likely to be perceived as separate entities. In our experiments, each continuity stimulus consists of 4 segments. Segments 1 and 3 share the same orientation and thus constitute a continuous line, and segments 2 and 4 share another orientation. The smaller the orientation difference between these two continuous lines, the more the entire stimulus follows a Gestalt principle (see Fig 9C in Methods).

We measured synchrony of firings of neurons in the model, in response to input stimuli. Importantly, we considered only those neurons which were receiving external input from the stimuli. The input images were used as a model input in the form of the Poisson spiking process. Please see Methods for a comprehensive description of every stimulus, and the procedure of transforming image stimuli into Poisson spiking input.

Our main measurement was Group Rsync. For every stimulus, we randomly selected 4 neurons from each stimulus segment and measured the synchrony of all their firings together (Fig 3A provides an example of selecting neurons for the measurement).

**Figure 3.**
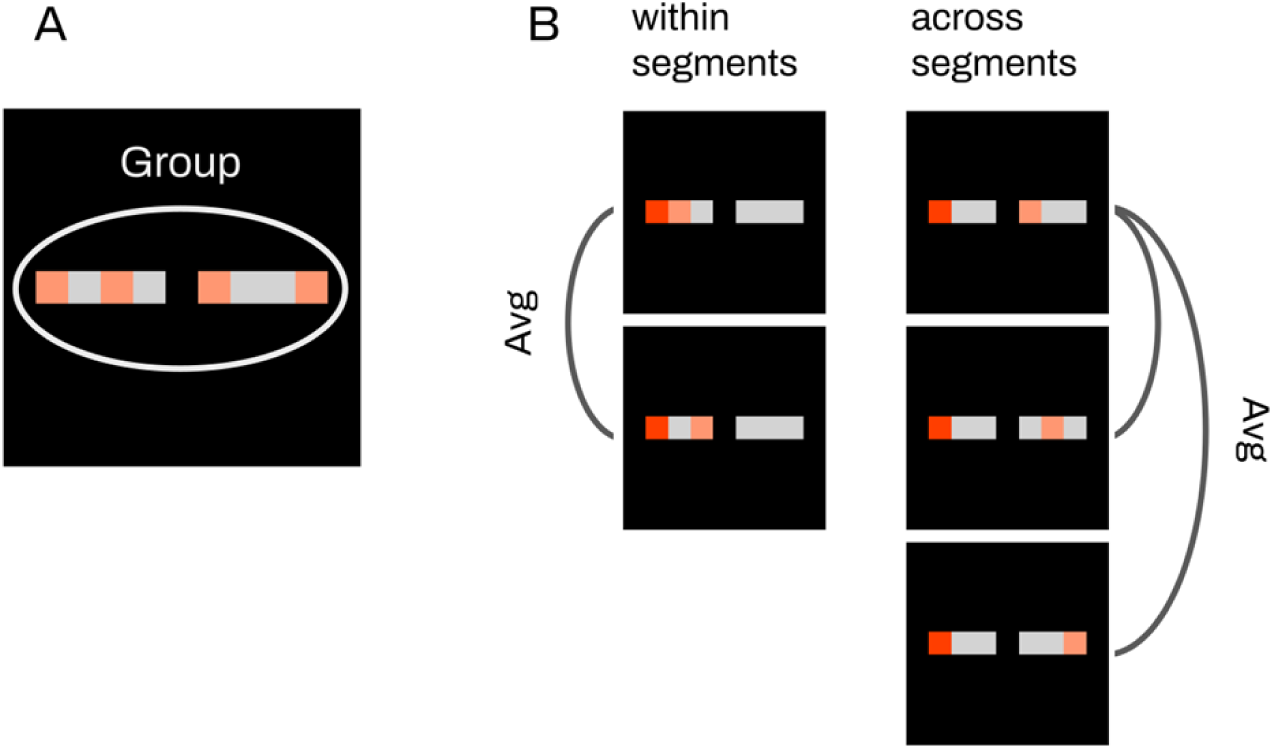
Selecting neurons for synchrony measurements. **A.** Group Rsync. Several neurons (2 in the example picture) are selected from each stimulus segment, then synchrony is measured for the group of all selected neurons. **B.** Avg Pairwise Rsync. For each neuron (consider the red neuron as an example), we first measured its firing synchrony with every other neuron from the same segment and averaged the result. Second, we measured its synchrony with every other neuron from the other segment and averaged this result too.

The amount of neurons to measure within one segment is arbitrary, we tested various numbers from 1 to 6 and did not observe any difference in the results.

To additionally illustrate the synchrony differences for various experimental conditions, we measured synchrony in a pairwise fashion (we further refer to it as Avg Pairwise Rsync): between all pairs of neurons from similar and different segments of the stimuli (see Fig 3B).

### Synchrony for minimal and maximal Gestalt-likeness

First of all, we measured the synchrony between firings of neurons receiving input from two different line segments. For the proximity and similarity principles, we considered left and right segments (Fig 4 A-B). For the continuity principle, the synchrony was measured between neurons which receive input from pairs of segments: left and top, left and right, right and bottom, top and bottom. From each segment, 5 neurons were randomly selected for synchrony measurements.

**Figure 4.**
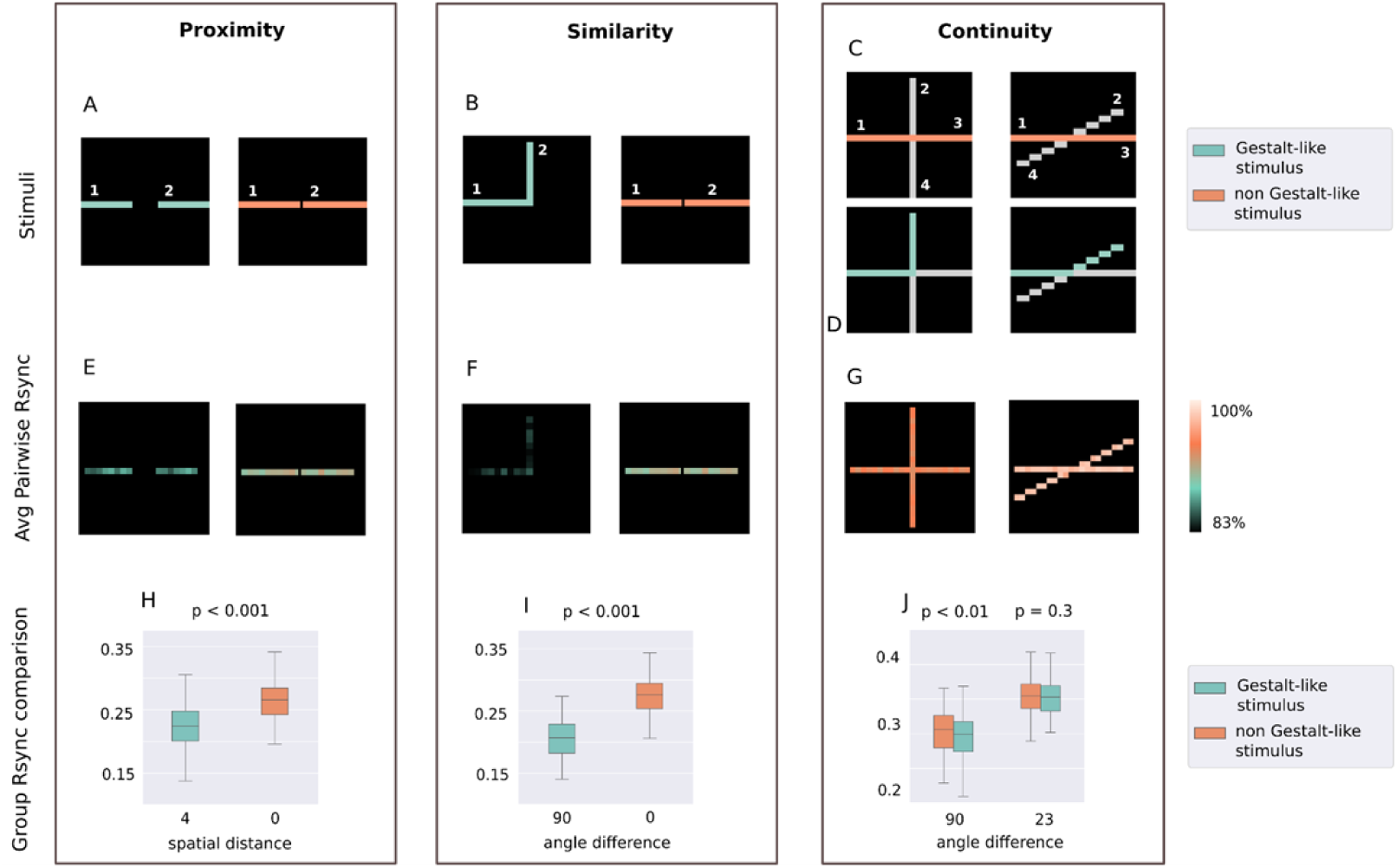
Rsync differences for most Gestalt-like and least Gestalt-like stimuli. **A-D.** Stimuli for proximity, similarity, and continuity are described in further detail in Methods. **E-G.** Average pairwise Rsync for each neuron’s spike train with every neuron from the other segment. Shown in the percentage from maximal measured Rsync. **H-J.** Group Rsync was measured within a group of 10 randomly selected neurons, 5 for each stimulus segment.

Measured synchrony reflected how closely the entire stimulus was following a certain Gestalt principle. Highly significant (p < 0.001) differences in synchrony were detected between stimuli with the greatest and the lowest Gestalt-likeness for the proximity and similarity principle, and significant difference (p < 0.01) for the continuity principle. We further report the results for all three principles.

For the proximity principle, the difference was measured as the size of the gap between two segments. The smaller the gap, the stronger the entire stimulus follows the proximity principle. Fig 4A represents stimuli under two conditions: minimal (0 pixels) and maximal (4 pixels) distance between two segments. The median synchrony was equal to 0.26 between two segments of the stimulus with a spatial distance equal to 0 (min. spatial distance). The median synchrony was equal to 0.22 for segments of the stimulus with the spatial distance equal to 4 (max. spatial distance). The non-parametric Wilcoxon rank test showed a significant difference with the p-value being less than 0.001.

For the similarity principle, the angle orientation difference between two line segments was measured. The smaller the difference, the closer the entire stimulus is following the Gestalt principle of similarity. Fig 4B represents stimuli under two conditions: minimal (0°) and maximal (90°) angle difference between two segments. The median synchrony was equal to 0.28 between two segments of the stimulus with an angle difference equal to 0° (both segments having equal angle orientation). For the stimulus with a between-segment angle difference equal to 90°, the median synchrony was only 0.2. The Wilcoxon rank test showed a significant difference with the p-value being less than 0.001.

For the continuity principle, the angle difference between two lines was measured, with a smaller difference indicating that the stimulus follows the continuity principle more strongly. Fig 4C represents stimuli under two conditions: minimal (23°) and maximal (90°) angle difference between two lines. For the largest angle difference (90°), the median synchrony between segments lying on the same line was equal to 0.306. The median synchrony between segments of different lines was equal to 0.299. The Wilcoxon rank test showed a significant difference with the p-value equal to 0.003. As for the smallest angle difference (23°), the median synchrony between segments of the different lines was equal to 0.353. The median synchrony between segments of the same line was equal to 0.355. The Wilcoxon rank test showed a non-significant difference with the p-value equal to 0.34. Thus, the difference in synchrony was more pronounced for a larger difference between the two lines. For the smallest difference (23°), the difference in synchrony was insignificant.

The difference in synchrony between more and less Gestalt-like stimuli was more prominent for the proximity and similarity experiments, compared to the continuity experiment. At the same time, the overall synchrony was greater for both continuity experiment setups (23° and 90° angle difference) than for any of the others. The maximal measured synchrony for the continuity experiments exceeded 0.4, while the maximal synchrony for proximity and similarity was under 0.35. Such inconsistency can be explained by the greater overall excitation in the continuity experiments, since more neurons were receiving both external input and lateral input from the horizontal connections. We assume that lateral input is crucial for establishing synchrony, since neurons have to to interact with each other in a recursive fashion through the synapses.

However, all the experiments with varying significance levels and effect sizes showed that the Gestalt-likeness of the input is reflected in the spiking synchrony of corresponding neurons.

### Group synchrony and the Gestalt structure of the stimulus

We tested how spike synchrony changes with respect to gradual changes in the Gestalt structure of the stimuli. For the Gestalt principles of proximity and similarity, synchrony increases gradually alongside the increasing Gestalt-likeness of the stimuli. For continuity, the difference in synchrony between Gestalt-like and non-Gestalt-like stimuli segments also increases simultaneously with the increasing difference between these segments.

In the proximity experiment, spatial distance between the stimulus segments was inversely proportional to the Gestalt-likeness of the stimulus. We used input stimuli with between-segment distances varied from 4 to 0 and measured group spike synchrony for each stimulus (Fig 5).

**Figure 5.**
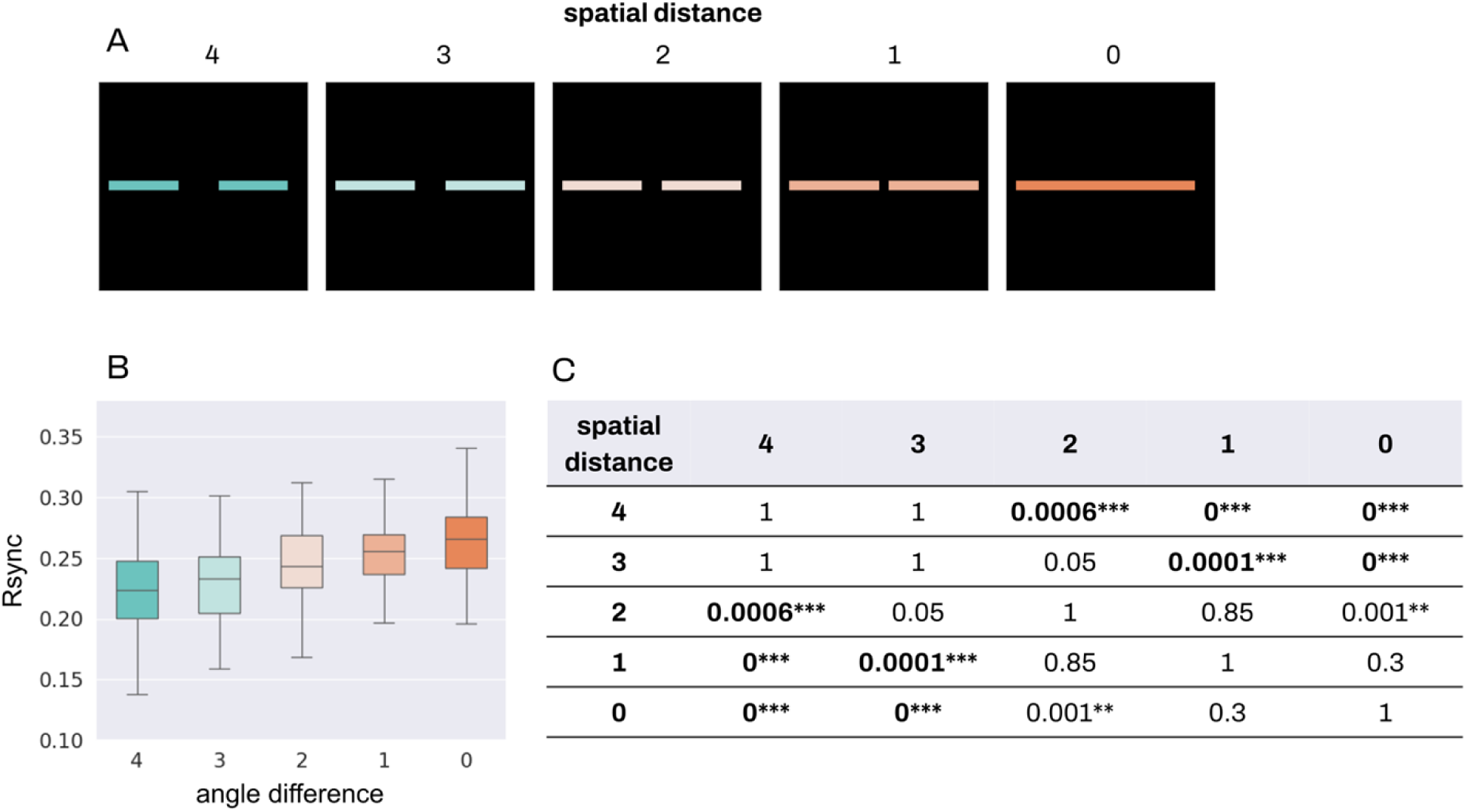
Group Rsync for proximity. Comparison of group Rsync for all stimuli conditions in the proximity experiment. For every stimulus, 4 neurons were randomly selected from each stimulus segment, and the group Rsync was measured between all 8 of them. **A.** Proximity stimuli with various distances between the segments. **B.** Comparison of group Rsync for all the stimuli. **C.** Results for non-parametric Dunn test with Bonferroni adjustment for multiple comparison, rounded to 4 decimal. * stands for p < 0.05, ** for p < 0.01, *** for p < 0.001.

Kruskal-Wallis test showed significant between-group differences with p-value < 0.0005, and the effect size equal to 0.22. We further applied nonparametric Dunn test with Bonferroni adjustment for multiple comparisons (Fig 4C) and found statistically significant differences between the following groups: distances 0 and 2 (p < 0.01), 0 and 3 (p < 0.001), 0 and 4 (p < 0.001); 1 and 3 (p < 0.001), 1 and 4 (p < 0.001); 2 and 4 (p < 0.001). Interestingly, differences for distances 1 and 2, 2 and 3, 3 and 4 were not significant. The most Gestalt-like stimulus with the 0 distance between segments led to significant differences in synchrony with the stimuli with all the other between-segment differences. The most significant differences were observed between stimuli with the greatest (0 distance) and lowest (3 and 4 distance) Gestalt-likeness.

In the similarity experiment, the angle difference between the stimulus segments was inversely proportional to the Gestalt-likeness of the stimulus. Similar to the proximity experiment, we used input stimuli with various angle differences from 90° to 0° and measured the group spike synchrony (Fig 6).

**Figure 6.**
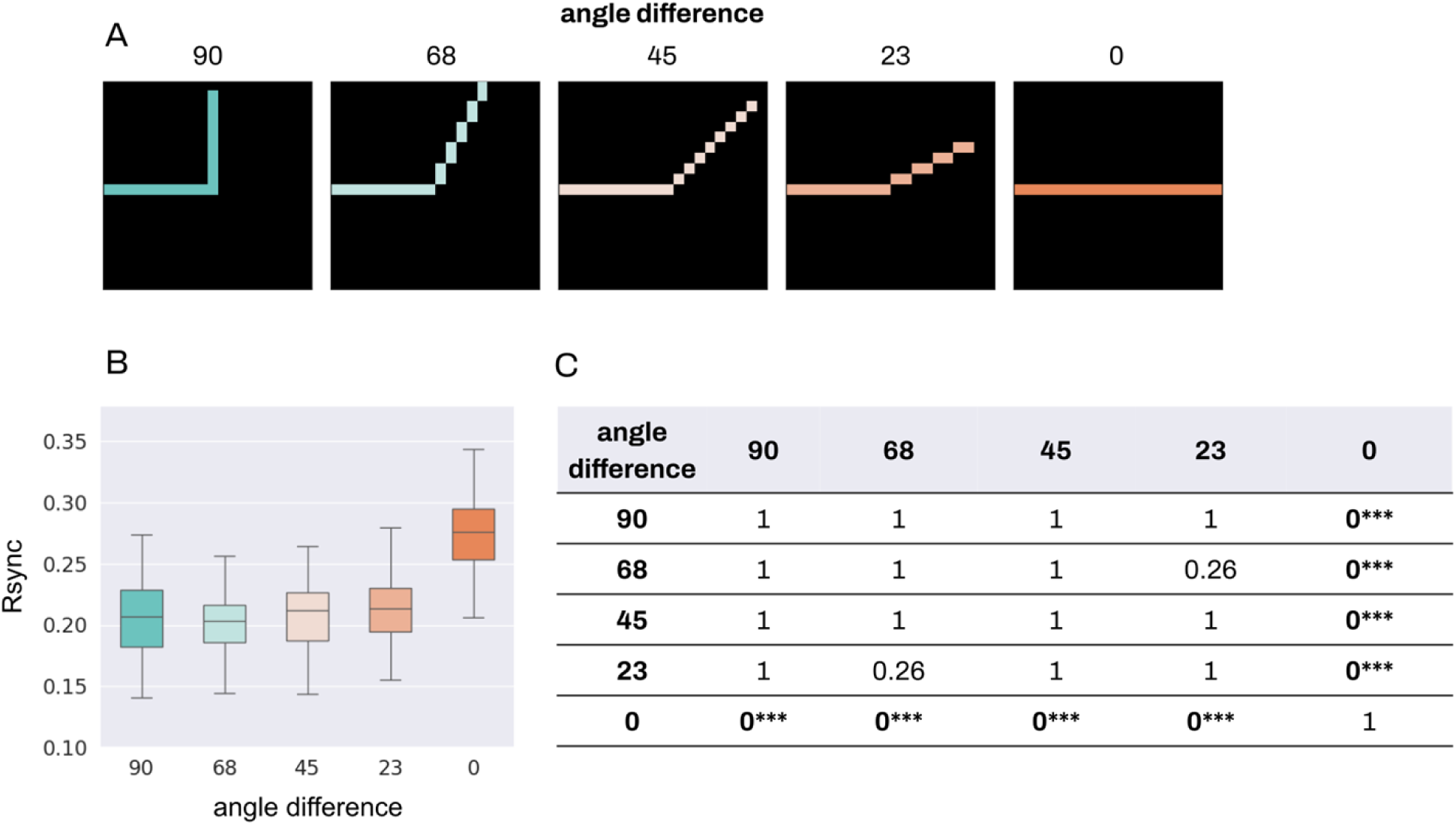
Group Rsync for similarity. Comparison of group Rsync for all stimuli conditions in the similarity experiment. For every stimulus, 5 neurons were randomly selected from each stimulus segment, and the group Rsync was measured between all 8 of them. **A.** Similarity stimuli with various angle differences between the segments. **B.** Comparison of group Rsync for all the stimuli. **C.** Results for non-parametric Dunn test with Bonferroni adjustment for multiple comparison, rounded to 2 decimal. *** stands for p < 0.001.

Kruskal-Wallis test showed significant between-group differences with p-value < 0.0005, and an effect size equal to 0.3. The subsequent non-parametric Dunn test with Bonferroni adjustment for multiple comparisons (Fig 6C) showed significant differences between the following groups: angle differences 0° and 23° (p < 0.001), 0° and 45° (p < 0.001), 0° and 68° (p < 0.001), 0° and 90° (p < 0.001). Only the most Gestalt-like stimulus with the 0° angle difference between segments led to highly significant differences in synchrony with the stimuli with all the other between-segment differences.

For each stimulus in continuity experiments, we always compared two groups: with stimulus segments constituting one line or lying on two different lines. The angle difference between lines varied from 90° to 23° (Fig 7A). Unlike similarity experiments, we did not consider a 0° angle difference for continuity, because in that case two lines would be merged into one, and segments 1 and 4, 2 and 3 would coincide. Since only two-group comparisons were conducted, we used a Wilcoxon rank test (Fig 7C).

**Figure 7.**
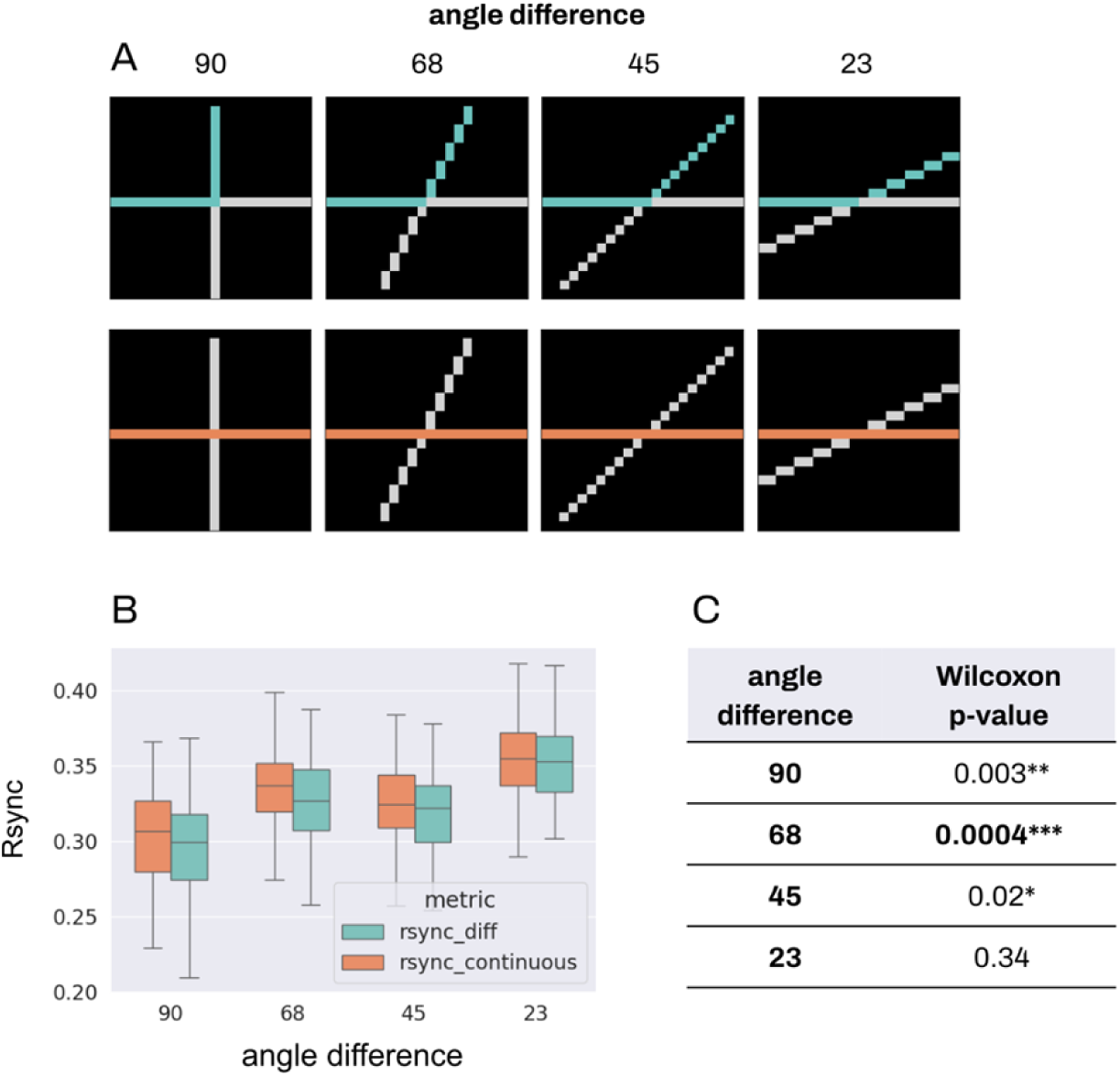
Group Rsync for continuity. Comparison of group Rsync for all stimuli conditions in the continuity experiment, for neurons from continuous and different segments of the stimuli. For every stimulus, 5 neurons were randomly selected from each stimulus segment, and the group Rsync was measured between all 10 of them. Please see Fig 9 in Methods for details on stimuli segmentation. **A.** Continuity stimuli with various angle differences between the segments. **B.** Comparison of group Rsync between different and continuous segments for all the stimuli. **C.** Results for Wilcoxon rank test, rounded to 4 decimal. * stands for p < 0.05, ** for p < 0.01, *** for p < 0.001.

For two smallest angle difference between lines and hence the greatest Gestalt-likeness (23°), we have not observed significant differences in synchrony. However, the greater angle difference and the lower Gestalt-likeness led to greater differences in synchrony. Significant differences were detected for greater angle differences: p < 0.05 for 45°, p < 0.001 for 68°, p < 0.01 for 90°. The 90° difference led to the second greatest significance level, and the results in general fell in line with our expectations: greater Gestalt-likeness corresponds to greater spike synchrony between segments.

There can be multiple reasons to explain the lower difference in synchrony between Gestalt-like and non-Gestalt combinations of segments (i.e. segments lying on the same or on different lines within one stimulus) for 90°. Since we measured the synchrony between segments 1 and 2, 3 and 4 (see Fig 9C for continuity stimuli segmentation), one can observe that spatial distance is lower between 1 and 2 for 90°, than for all other angles. Thus, while the angle difference between segments increases, the spatial distance decreases at the same time. Both spatial distance and orientation similarity contribute equally to the interneuronal connectivity strength, which we assume is the basis for establishing synchrony between neuron firings. Also, the continuity stimuli are more complex than similarity stimuli, despite the similar variety at the angle difference. Thus, the model receives more overall excitation, which may lead to increase in the system’s response variability and unexpected results.

However, it is worth noting that the result of the Wilcoxon test still remains significant in this case. Also, we do not observe a significant difference in synchrony for the most Gestalt-like stimulus with the 23° angle difference, despite the greatest spatial distance between neurons. These two observations lead to the conclusion that synchrony still reflects the Gestalt structure of the continuity stimuli, although the effect is not as pronounced as for the similarity experiment.

We also emphasize that the synchrony differences in response to various Gestalt likeness were measured for the entire group of activated neurons, which means that the information about the stimulus is shared across all neurons receiving input. Thus, synchrony can operate as an emergent phenomenon encoding global information about the entire stimulus.

### Average pairwise synchrony and the Gestalt structure

Fig 8 illustrates the difference between the Avg Pairwise Rsync within similar and different line segments. For all three types of experiments (proximity, similarity, and continuity), Rsync within one stimulus segment was higher than between neurons belonging to different segments. At the same time, Rsync both within one segment and between two segments was increasing in accordance with the Gestalt-likeness of the stimulus. Thus, the smaller the distance between two segments for proximity stimuli (Fig 8A), the greater both within- and between-Rsync was. For the similarity and continuity stimuli, the smaller the angle difference between two segments, the higher the pairwise Rsync between them (Fig 8 B-C). Thus, the more similar the stimuli segments were to each other – the higher spike synchrony was observed between neurons tuned to different segments of these stimuli.

**Figure 8.**
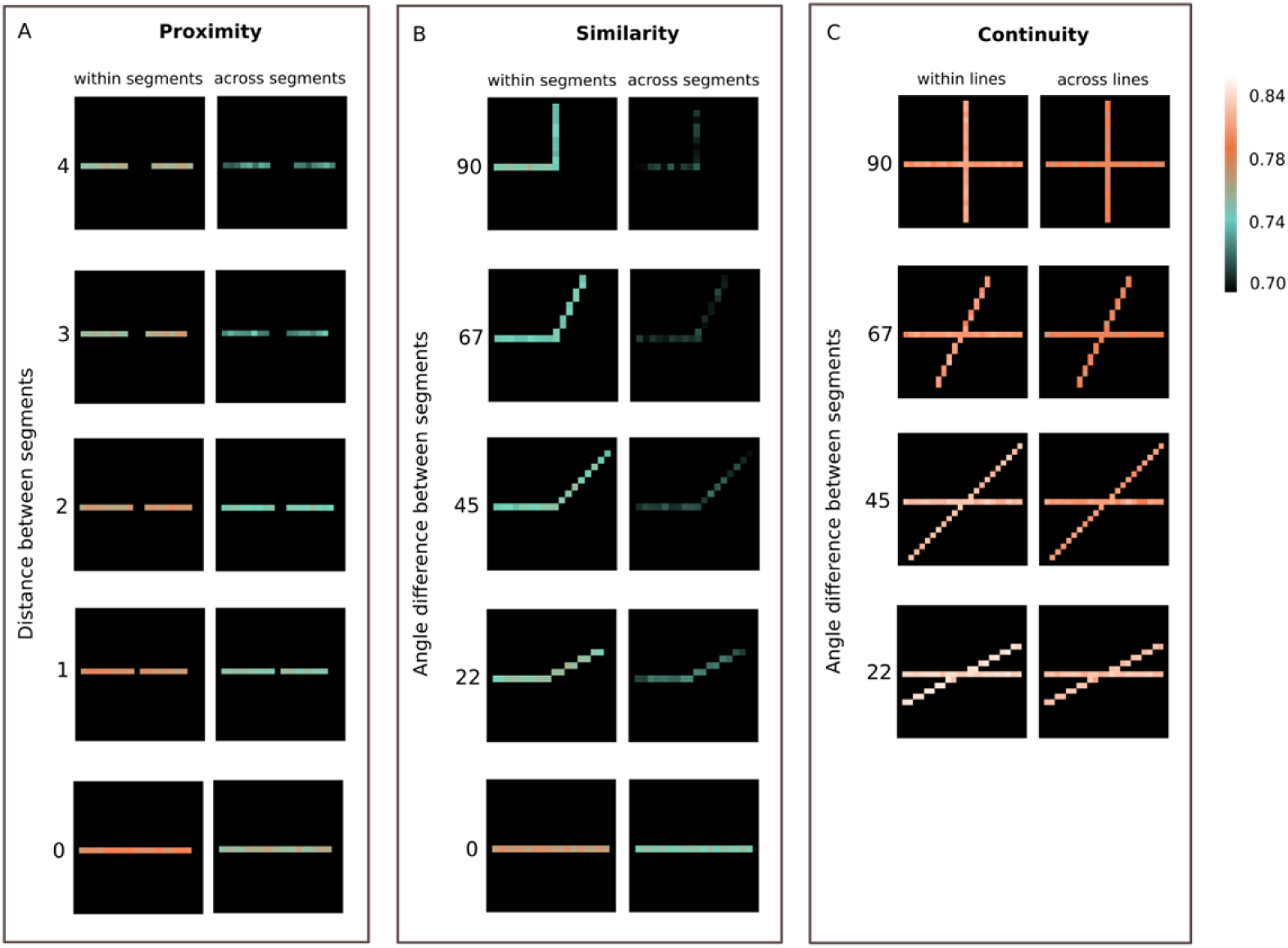
Avg Pairwise Rsync. Avg Pairwise Rsync measured for each stimulus separately. Only neurons receiving the external input are shown. Rsync of each neuron’s spike train with every neuron from the similar and the other segment is measured pairwise and then averaged. The corresponding stimuli with the segmentation are presented in Fig 9 A. **A-B.** Avg Pairwise Rsync measured for each stimulus separately, for proximity and similarity experiments. On the left, for each neuron the synchrony with the other neurons of its segment is measured. On the right, for each neuron of segment 1 the synchrony with the neurons of segment 2 is shown, and for each neuron of segment 2 the synchrony with the neurons of segment 1 is shown. **A.** Avg Pairwise Rsync is demonstrated for stimuli with various distances between the segments. **B.** Avg Pairwise Rsync is demonstrated for stimuli with various angle differences between the segments. **C.** Avg Pairwise Rsync measured for each stimulus separately, for continuity experiments. On the left, for each neuron of segment 1 the synchrony with the neurons of segment 3 is shown, and for each neuron of segment 3 the synchrony with the neurons of segment 1 is shown. On the right, for each neuron of segment 1 the synchrony with the neurons of segment 2 is shown, and for each neuron of segment 3 the synchrony with the neurons of segment 4 is shown. Mean pairwise synchrony is demonstrated for stimuli with various angle differences between the lines.

This illustrates that spike synchrony can be measured not only in a group, but also in a pairwise fashion, and still be an indicator of how closely the stimulus is following the Gestalt principles. Since the pairs for measurement included neurons within the stimulus on various distances from each other, this stays in line with the argument about the global nature of spike synchrony. It should also be noted that overall Avg Pairwise Rsync stays maximal for continuity experiments. Again, this observation is consistent with the Group Rsync increasing for continuity experiments due to the greater amount of both external and lateral excitatory input to neurons.

Additionally, we investigated how spike synchrony depends on the connectivity structure in the network, since synchrony in V1 was shown to arise from the horizontal intracortical connections. Avg Pairwise Rsync appeared to be correlated with the connectivity strength between neurons, with the Pearson r correlation coefficient up to 0.42. It shows that the connectivity structure is reflected in the patterns of synchronous firing to a certain extent. For more detail on correlation analysis please see Supplementary materials.

## Discussion

We showed that fully-excitatory spiking networks in a noisy environment demonstrate synchronous response to the Gestalt-like visual stimuli. The better stimuli follow the said Gestalt principles (proximity, continuity and similarity), the greater is spike synchrony. This demonstrates the computational role of spike synchrony as a mechanism for estimating the match between the stimulus and a prior encoded in a network connectivity structure.

Often co-activated cortical cells are well connected, which reflects the long-term visual experience. We argue that synchrony between such strongly connected units can serve as a measure of how well the perceived stimulus matches the acquired visual experience. Thus, its increase in response to Gestalt-like stimuli (unbroken lines, lines with the segments of similar orientation, etc.) shows that most often-experienced behaviorally relevant objects in the visual environment tend to follow the Gestalt structure. This is consistent with the natural statistics observations of [41, 45–48]. That is, if the horizontal connections reflect the most often experienced visual objects and serve the basis for spike synchrony, it should be natural for neurons to synchronize in response to Gestalt-like stimuli.

From the Bayesian perspective, we demonstrated that synchrony in the model of V1 can be interpreted as a familiarity of an encoded stimulus, which is a match of prior probability of an input that is encoded in the network structure, and the stimulus that is encoded in the local spike rate of neurons [15]. This mechanism is consistent with the observed spiking data recorded under stimuli conditions reflecting the Gestalt law [1].

The transition of a strictly local encoding of stimulus features, towards an emergent property that forms spatially extended representation based on spike synchrony, is a crucial and efficient computational feature. It allows separate processing of the stimulus content on the local level, and processing based on global properties linked to stimulus familiarity. This can be, for example, a proxy for how well a stimulus is encoded (low familiarity corresponds to weak information and vice versa), or whether the stimulus is new or has been encountered before. For agents, animals and humans, this information might be at least as relevant as the stimulus information itself. This emergent proxy will allow the agent to plan and perform continuous actions that can be stopped if the collected information is sufficient. Such active sampling and inference is discussed in different aspects across the fields of cognitive science, neuroscience and machine learning, and seems to be best described by the theory of Posterior sampling in reinforcement learning [52]. We therefore believe that spike synchrony, as an emergent phenomenon of spiking neural networks with strictly local stimulus feature encoding, is a computational principle that allows for the inference of global features, i.e. stimulus family, that can be relevant for efficient active learning and sensing.

## Methods

### Input stimuli

In our study, we examined the relationship of spike synchrony and the binding of visual stimulus components for three Gestalt grouping cues: proximity, similarity, and continuity [42]. The principles can also interact with each other, either strengthening or weakening their effects when combined.

1. Proximity principle: According to Gestalt theory, the principle of proximity states that when single visual components are located near each other, they are perceived as forming a larger visual composition. The aggregation of individual elements is grouped together to create a new superordinate entity.
2. Similarity principle: Another principle of Gestalt theory is the similarity principle, which involves integrating visual components into perceptual groups based on the similarity of their appearance. This principle classifies elements according to their visual features.
3. Continuity principle: The continuity principle, a fundamental aspect of Gestalt theory, organizes patterns into new compositions based on the alignment of their established direction. Conversely, patterns with abrupt changes in orientation are partitioned into distinct elements. For intersecting lines, the concept of Gestalt-likeness relates to how effectively the lines form a continuous shape or pattern. The continuity principle helps distinguish stimuli with visual overlap.

For the experiments, we ran simulations with a spiking neural network receiving artificial stimuli. The visual input for our network was generated from simple artificial images of size 20 by 20 pixels. These stimuli represent the input for the early stages of the visual pathway, constituting simple cells recognizing edges with location and orientation selectivity and implementing the examples of Gestalt theory principles (1) - (3) respectively. Before being presented to the network, input stimuli are partitioned into different channels by an edge detector sensitive to orientation. This reflects the selectivity of the visual cortex to edges of certain angles through hypercolumns and orientation columns. Edges in the input stimuli are undirected, limiting the relevant range of orientations from 0° to 180°. Since the size of our networks grows exponentially with the number of orientation channels, for reasons of computational efficiency, in the experiments only input stimuli with an orientation between 0° to 90° were considered. Sensitivity to the degree of orientation was set to 22.5°, resulting in 5 channels for input stimuli of orientations [0°, 90°], see Fig 8. The network receives pixel-wise probabilistic input spikes from the stimulus at a rate of 40Hz with a temporal resolution of 0.5ms. Probabilistic input is based on a Poisson-point process — a mathematical model for generating random events that occur independently of each other over time with a constant average rate of occurrence and is often used to capture the stochastic nature of neural activity.

Additionally, we introduced Gaussian noise at every time step for random variability into the input, simulating real-world conditions where input signals are subject to fluctuations. The signal-to-noise ratio was set to 0.4, with any higher ratio resulting in the onset of runaway excitation. During runaway excitation, the activity within the network is only limited by the spike-rate saturation leading to the constant firing of neurons without any meaningful modulation or information processing.

A new set of stimuli patterns was created for each experiment, each reflecting a range of Gestalt-likeness of the experiment’s Gestalt principle respectively. The effective representation of the range of degrees in Gestalt-likeness closely follows the definition of Gestalt principles, with the assumption that our stimuli models similar patterns present in the early processing of the visual pathway sufficiently close.

Input stimuli for the Gestalt principle of proximity are based on the spatial distance between segments (see Fig 9A). The closer the two segments are, the more likely they will be perceived as a group. As the distance between segments increases, so does the likelihood of perceiving them as separate entities. Therefore, the degree of Gestalt-likeness of stimuli decreases with an increase of range between two segments. Figure 9A shows a set of 5 stimuli with an interval in pixel distances between segments from 0 to 5 which was used in the proximity experiments.

**Figure 9.**
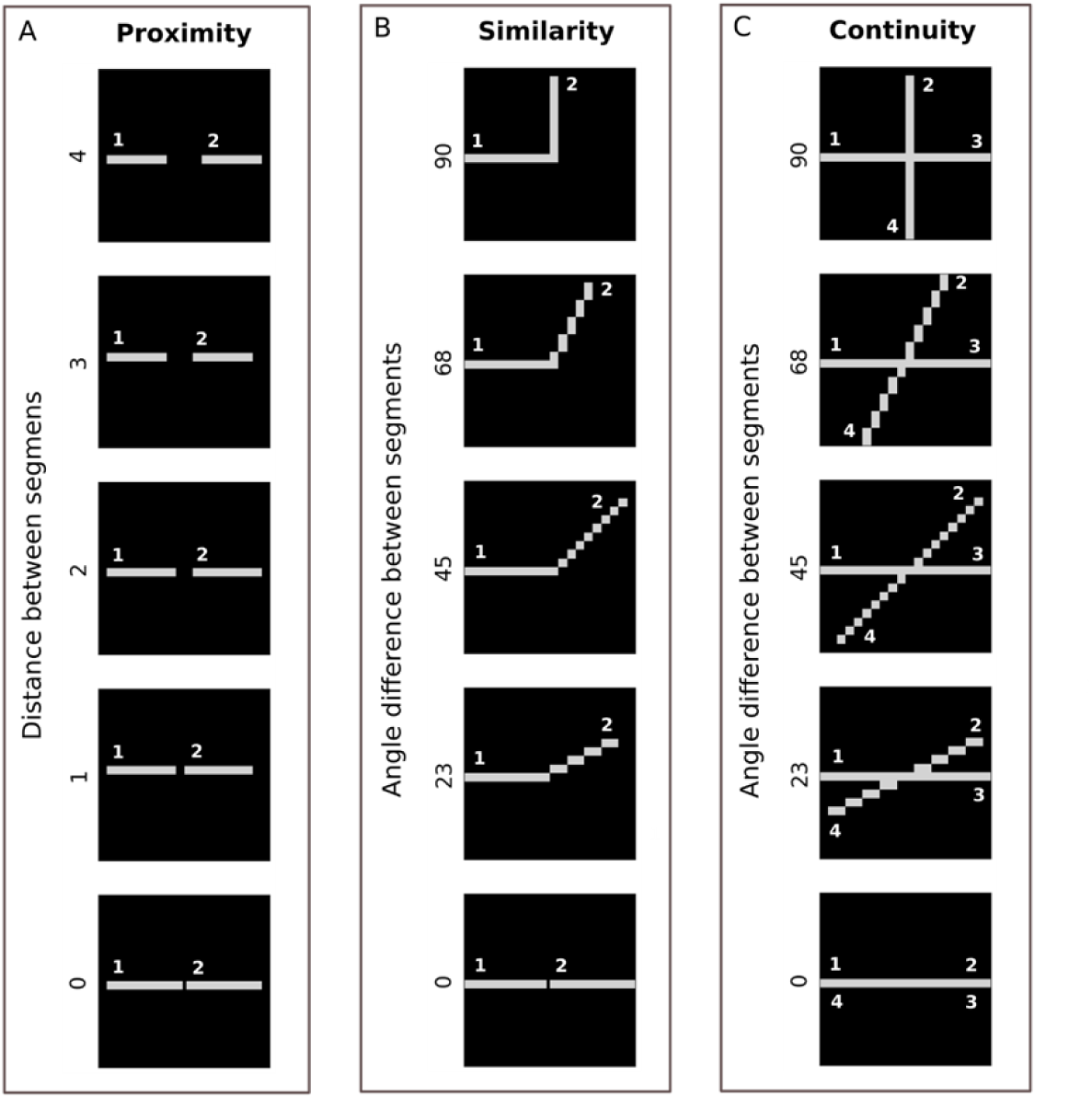
Input stimuli. Input stimuli for experiments in descending order of their adherence to Gestalt-likeness. **A.** Input stimuli for the Gestalt principle of proximity. Each stimulus consists of two segments: 1 and 2. The spatial difference between segments varies from 4 to 0. **B.** Input stimuli for the Gestalt principle of similarity. Each stimulus consists of two segments: 1 and 2. The angle orientation difference between segments varies from 90° to 0°. **C.** Input stimuli for the Gestalt principle of continuity. Each stimulus consists of four segments: 1, 2, 3, 4. Segments 1 and 3, 2 and 4 constitute continuous lines. The angle difference between lines varies from 90° to 0°.

Input stimuli for the Gestalt principle of similarity are based on the orientation similarity, the similarity of segments’ degree of rotation (see Fig 9B). The closer two segments are in visual appearance the higher their Gestalt-likeness. Segments with minimal orientation differences are perceived as having high orientation similarity and are more likely to be perceived as a group. As the difference in orientation between segments increases, making them visually more distinct, so does the likelihood of perceiving them as separate segments. Therefore, the degree of Gestalt-likeness of stimuli decreases. Figure 9B shows a set of 5 stimuli with 90°, 68°, 45°, 23° and 0° orientation of the second stimulus segment, increasing in their Gestalt-likeness, that were used in the similarity experiments.

Input stimuli for the Gestalt principle of continuity are based on the abruptness of a segment’s change of direction in two intersecting lines (see Fig 9C). Elements arranged in a continuous manner are more likely to be perceived as belonging together. Segments with minimal orientation differences are perceived as having high orientation similarity and are more likely to be perceived as a continuous line. Additionally, as the difference in orientation between two lines decreases so does the likelihood of perceiving the line’s segments as a disjoint part that belongs to the other’s segment instead. Therefore, a smaller difference indicates that the stimulus follows the continuity principle more strongly. Figure 9C shows a set of 5 stimuli with 90°, 68°, 45°, 23° and 0° orientation increasing in their Gestalt-likeness. Unlike in the similarity experiments, a 0° angle difference for continuity was not considered for the experiments, since here both lines fully merge into one, and segments 1 and 4, 2 and 3 coincide.

### Model

To simulate neurons, we used the Izhikevich model for spiking neural networks [51] with excitatory pulsed coupling. The Izhikevich model of a spiking neuron emulates the dynamics of the classical Hodgkin-Huxley model [53] but operates more efficiently. It is a two-dimensional system, where each neuron is characterized by two internal variables: the membrane potential v_i_ and the recovery variable u_i_. In our network only excitatory connections are used.

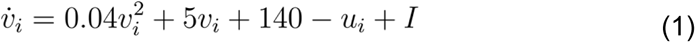

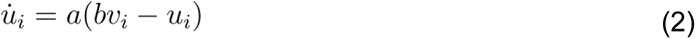

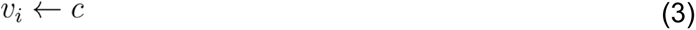

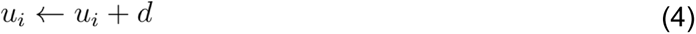

The equations 1 and 2 describe how the values of v and u change over time, and i denotes the time moment. In equation 2, a stands for the time scale of recovery u. The bigger a leads to faster recovery. In our model, we consider a = 0.02. The parameter b describes the sensitivity of the recovery variable to the fluctuations of the membrane potential v. We set b = 0.2. When the value of v exceeds the activation threshold 30 mV, the neuron is considered activated and we record a spike event. After the spike is detected, the values of v and u are being reset according to equations 3 and 4. The voltage reset constant c is set to -65 mV, and the recovery update variable d is set to 6.

The parameter I in equation 1 defines the entire input to the neuron. It consists of input currents received from lateral connections (I^net^) and the external stimulus (I^up^). The external input is sent to the network in a probabilistic form, as a Poisson process with an average firing rate of 40 Hz (see Methods. Input stimuli for details).

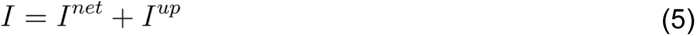

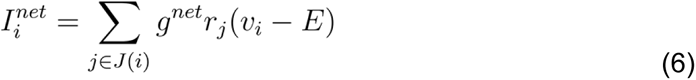

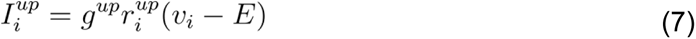

Parameter g represents the upper limit of the lateral (g^net^) and the upstream (g^up^) synaptic conductances. The synaptic reversal potential E is set to 0. The conductance of each synaptic connection is adjusted by the open receptor fraction r, influenced by incoming spikes.

For both upstream and lateral connections, r is driven by neurotransmitter concentration in the synaptic cleft [T]_j_, which in turn is represented by a pulse of duration τ = 0.02 after each incoming spike.

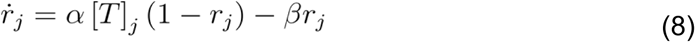

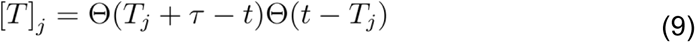

r_i_ is parameterized, also for both upstream and lateral connections, by the rise and decay time constants α = β = 8. The transmitter concentration [T]_j_ is the product of two heaviside step functions Θ. They define that neurotransmitter is present in the synaptic cleft ([T]_j_ = 1) starting from the occurrence of a presynaptic spike at the moment T_j_, until the moment T_j_ + τ.

### Connectivity

A network architecture was designed to reflect the retinotopic organization of the visual cortex with artificial spiking neurons. It comprises a single n×m layer of Izhikevich neurons where n, m correspond to the stimulus dimensions. The fixed connectivity of the network structure models the retinotopic organization of the visual cortex (V1) and is organized as hypercolumns of neurons selective towards the location and the orientation of stimuli. Specifically, the connection strength between neurons is determined by two factors: the spatial proximity of the neurons’ receptive fields and the similarity in the angle orientation of the stimuli (Fig 10).

1. Spatial proximity: Each neuron is receptive to location-specific input from both external stimuli matching their retinal coordinates and horizontal connections by all other neurons with their connection strength inversely proportional to their distance. In this way, neighboring neurons with similar receptive fields will receive stronger input than distant ones of dissimilar receptive fields. As a distance metric, we used the Chebyshev distance, which quantifies the maximum difference between corresponding components of two vectors in a multi-dimensional space. The distance between two points, A(x1, y1) and B(x2, y2), in a two-dimensional space is here defined as the maximum absolute difference between their corresponding coordinates along the x and y axes.
2. Orientation similarity: Furthermore, neurons within the network are also selective to the orientation of stimuli, which is manifested in the formation of orientation-selective cortical columns. A subnetwork of neurons that receive input from the same region but select for different orientations represents an orientation-selective hypercolumn. All neurons in the network are connected to each other, with a connection strength that is inversely proportional to the angle difference.

**Figure 10.**
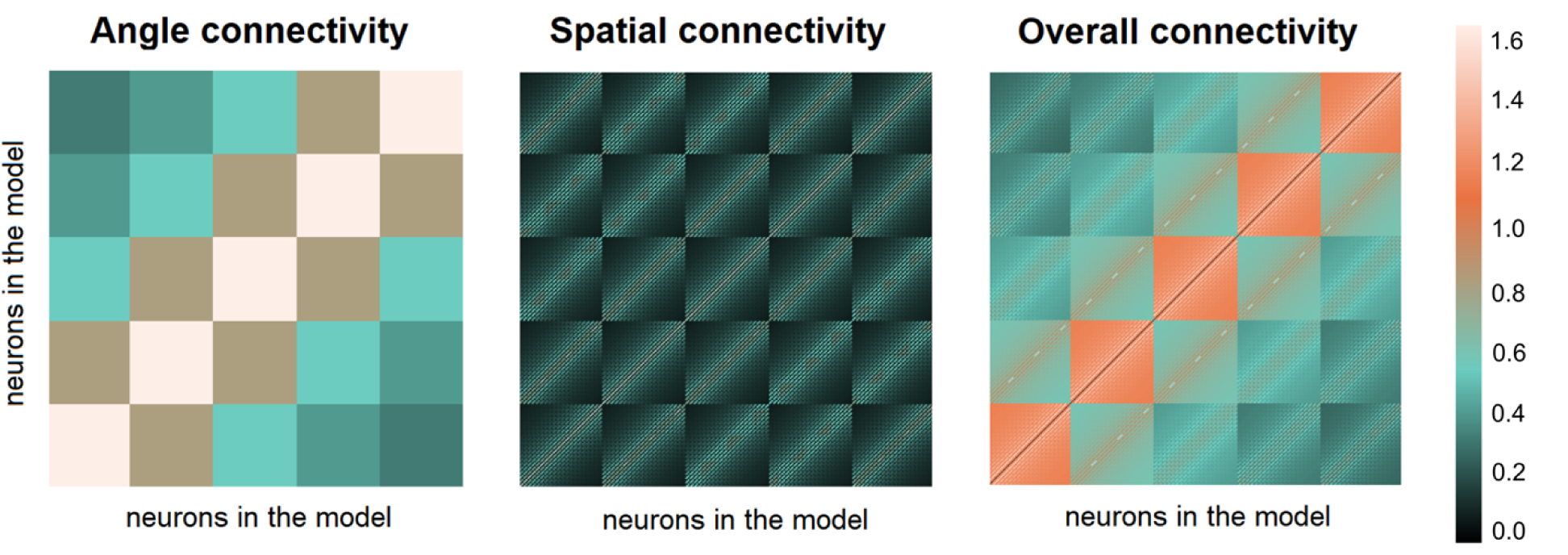
Connectivity Matrix. **Angle connectivity** defines the connection strength solely depending on the orientation similarity of neurons. The connection strength is inversely proportional to the angle difference. **Spatial connectivity** defines the connection strength depending on spatial distance between neurons. The connection strength is inversely proportional to their Chebyshev distance. **Overall connectivity** matrix is constructed as a sum of angle and spatial connectivity matrix values.

By taking both spatial distance and orientation similarity into account for building a connectivity matrix, our proposed network model emulates a simple retinotopic organization of V1 and provides a mechanism for the representation of location and orientation-specific stimuli through the connectivity of neurons within the network.

Importantly, spatial proximity and orientation similarity have an equal impact on the overall connection strength. No self-connections or inhibitory connections were used in our network. A detailed list of all hyper-parameters can be found in Appendix.

### Measure synchrony

We measure the temporal synchrony of the neuron spike traces following the procedure from Korndörfer et al. 2017 by adopting the Rsync metric (Eq. 10) [15]. It computes the average degree of zero-lag synchrony of the network, which ranges from 0 with all neurons firing out of phase, to 1 with all neurons strictly simultaneously firing.

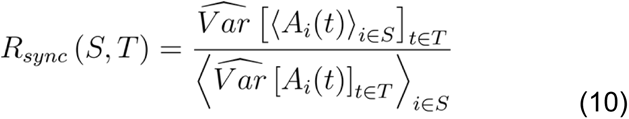

Here, A_i_ is an activation trace of the neuron i from the population S. To retrieve this activation trace, the raw binary spike train is first convolved with a causal exponential kernel k(t) = e^-2t^. Then, synchrony of a neuron population S is measured on a time interval T. It is given by the variance of the mean activation trace of this population, divided by the average variance of all neurons in the population. Intuitively, when all neurons are perfectly synchronous, i.e. fire precisely at the same time, the variance of their average activation trace is equal to a variance of each individual neuron and, respectively, to their average variance.

In the synchronized case, the sum of signals will display large-amplitude oscillations, while in the unsynchronized case, the individual signals will be out of step with each other and their sum will be nearly constant. Subpopulations of the network are drawn from neurons that receive external input from the stimulus segment, either from the same stimulus segment for average within-stimulus synchrony or across stimulus segments for average between-stimulus synchrony.

## Supplementary

### Avg pairwise Rsync reflects the connectivity structure

In V1, spike synchrony arises from the horizontal intracortical connections [26–28]. To show how it depends on the connectivity structure in the model, we visualized matrices of connections between neurons and pairwise synchrony matrices next to each other (Fig 11). Only neurons receiving the external input were considered.

**Figure 11.**
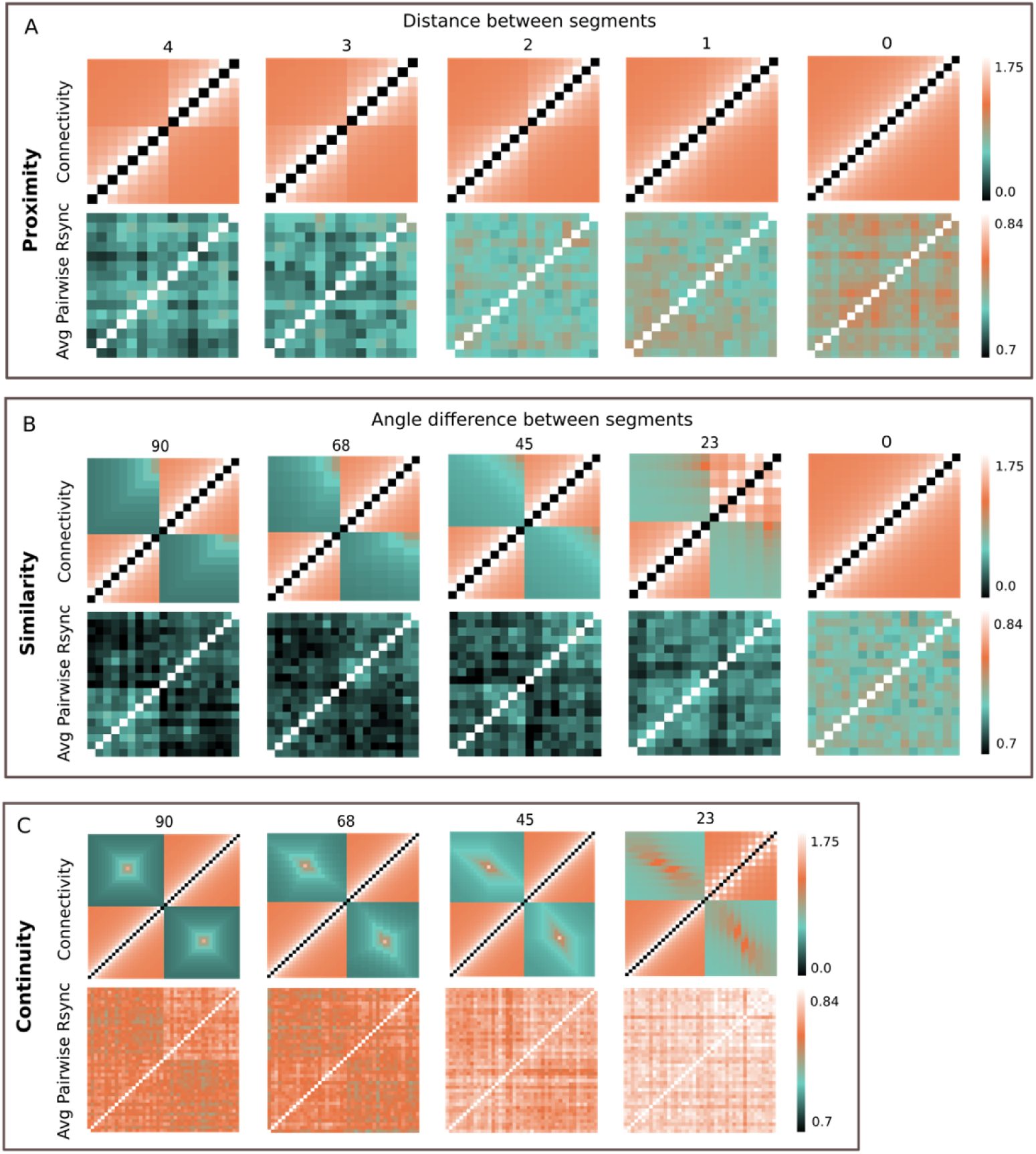
Rsync & connectivity. Comparison of the connectivity matrix and Avg Pairwise Rsync values for a group of neurons receiving the external input. **A.** Connectivity matrices and corresponding Avg Pairwise Rsync maps for proximity experiments. **B.** Connectivity matrices and corresponding Avg Pairwise Rsync maps for similarity experiments. **C.** Connectivity matrices and corresponding Avg Pairwise Rsync maps for continuity experiments.

What we found is that the connectivity structure was to a certain degree reflected in Avg Pairwise Rsync values. First, in all experiments the higher overall connectivity strength was leading to higher synchrony. Second, the structure of the connectivity matrix was reflected in the structure of the pairwise synchrony matrix to a certain degree. The structure of the synchrony matrix was able to capture the most pronounced structural details of the connectivity matrix. For example, the alterations of high and low connectivity values for proximity weight matrices were also visible in the corresponding pairwise synchrony matrices.

To quantify the similarity between connectivity and Avg Pairwise Rsync, we measured the cross-correlation between two matrices for all experimental conditions, averaged over 100 trials each (Table 1).

**Table 1.** Correlation between connectivity & pairwise Rsync. Pearson r correlation coefficients between Connectivity and Avg Pairwise Rsync, rounded to 2 decimal. P-values are < 0.005 for all experiments.

| Proximity |  | Similarity |  | Continuity |  |
| --- | --- | --- | --- | --- | --- |
| distance | Pearson $r \pm \text{sd}$ | angle difference | Pearson $r \pm \text{sd}$ | angle difference | Pearson $r \pm \text{sd}$ |
| 4 | $0.41 \pm 0.04$ | 90° | $0.33 \pm 0.06$ | 90° | $0.24 \pm 0.07$ |
| 3 | $0.39 \pm 0.05$ | 68° | $0.32 \pm 0.02$ | 68° | $0.23 \pm 0.06$ |
| 2 | $0.39 \pm 0.04$ | 45° | $0.32 \pm 0.07$ | 45° | $0.24 \pm 0.12$ |
| 1 | $0.39 \pm 0.04$ | 23° | $0.33 \pm 0.04$ | 23° | $0.2 \pm 0.1$ |
| 0 | $0.37 \pm 0.04$ | 0° | $0.38 \pm 0.04$ | 0° | |

What we found is that measured Pearson r values exceeded 0.3 in most cases, which implies a positive correlation. The correlation was the strongest for proximity experiments, likely due to neurons always sharing the same angle orientation and thus having a more homogeneous connection weights distribution. In contrast, the connectivity in continuity experiments tends to form more complex patterns, due to the more complex stimuli. It seemed to influence the correlation strength, which was lower than the average correlation strength for proximity and similarity experiments. For all the experiments spike synchrony was moderately or weakly correlated with the interneuronal connectivity structure.

### Group synchrony of time-jittered spike trains

Throughout the experiments we hypothesized that spike synchrony provides an additional information crucial for recognizing stimulus familiarity, and it highly relies on the temporal structure of the spike trains. To test whether the spike synchrony really depends on this fine temporal structure and not on e.g. firing rate, we did a control measurement for all three experiments with group synchrony. For each spike train, spike times were randomly jittered across the time scale: each spike time was randomly shifted in time within the range of (-80; +80) ms from its actual spiking. Then Rsync was measured on such modified spike trains with destroyed temporal structure (Fig 12).

**Figure 12.**
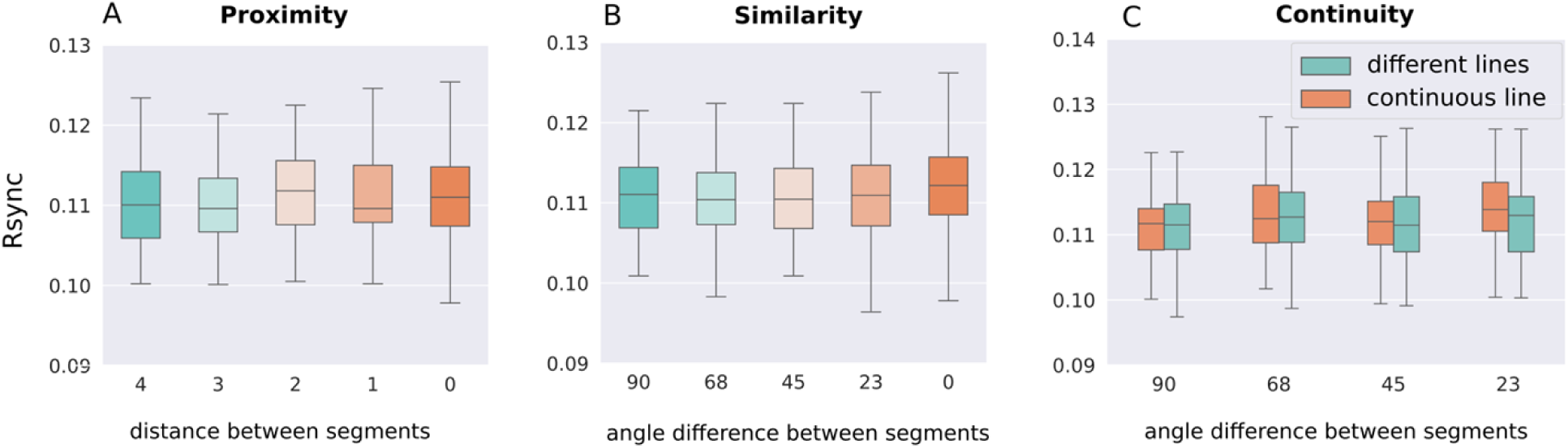
Group Rsync between segments for time-jittered spikes. Comparison of group Rsync for all stimuli conditions on time-jittered spikes. For every stimulus, 5 neurons were randomly selected from each stimulus segment, and the group Rsync was measured between all 10 of them. **A.** Comparison of group Rsync for proximity stimuli with various distances between the segments. **B.** Comparison of group Rsync for similarity stimuli with various angle differences between the segments. **C.** Comparison of group Rsync for continuity stimuli with various angle differences between the lines constituting a stimulus.

In contrast with the original experiments with non-jittered spike trains (see Results), for jittered spike trains synchrony does not change depending on the Gestalt structure of the stimulus. Not only no significant differences between experimental conditions are observed, but also the overall synchrony level is much lower than for the original non-jittered spike data. This holds true for all three Gestalt experiments. Thus, the temporal structure of spike trains contains synchrony information which is crucial for detecting Gestalt structure and hence, in our setting, prior probability of the input stimulus.

We applied similar statistical tests to time-jittered spike trains, as to the original spike data. For the proximity experiments, Kruskal-Wallis test showed a non-significant result with the effect size = 0.005 and p-value = 0.17, while the same test on the non-jittered data revealed significant between-group differences with the effect size = 0.16 and p-value < 0.001. We further applied a nonparametric Dunn test with Bonferroni adjustment for multiple comparisons (Table 2). It found no significant differences in synchrony between any stimuli.

**Table 2.** Dunn test for proximity experiments with time-jittered spike trains. The results of Dunn test with Bonferroni correction for groups with various spatial distances between stimulus segments for time-jittered spike trains, rounded to 2 decimal.

| spatial distance | 0 | 1 | 2 | 3 | 4 |
| --- | --- | --- | --- | --- | --- |
| 0 | 1 | 1 | 1 | 1 | 1 |
| 1 | 1 | 1 | 1 | 1 | 1 |
| 2 | 1 | 1 | 1 | 0.43 | 0.41 |
| 3 | 1 | 1 | 0.43 | 1 | 1 |
| 4 | 1 | 1 | 0.41 | 1 | 1 |

For time-jittered data from the similarity experiments, Kruskal-Wallis test did not find any significant between-group differences with the effect size = 0.007 and p-value = 0.12, whereas the effect size = 0.38 and p-value < 0.001 for non-jittered data. The subsequent non-parametric Dunn test on time-jittered data confirmed the absence of any differences in synchrony between stimuli with various angle differences (Table 3), although for the original non-jittered data Kruskall-Wallis test with Dunn post hoc test revealed multiple significant differences.

**Table 3.** Dunn test for similarity experiments with time-jittered spike trains. The results of Dunn test with Bonferroni correction for groups with various angle differences between stimulus segments for time-jittered spike trains.

| angle difference | 0 | 23 | 45 | 68 | 90 |
| --- | --- | --- | --- | --- | --- |
| 0 | 1 | 0.66 | 0.17 | 0.33 | 0.36 |
| 23 | 0.66 | 1 | 1 | 1 | 1 |
| 45 | 0.17 | 1 | 1 | 1 | 1 |
| 68 | 0.33 | 1 | 1 | 1 | 1 |
| 90 | 0.36 | 1 | 1 | 1 | 1 |

For continuity experiments with time jittering, we again followed the procedure for the original spiking data and compared two groups: with stimulus segments constituting one or two different lines. The angle difference between lines varied from 90° to 23°. We applied a Wilcoxon rank test and have only revealed a slightly significant result for the difference 23° (Table 4), unlike when measuring synchrony on the original data.

**Table 4.** Wilcoxon rank test for continuity experiments with time-jittered spike trains. The results of the Wilcoxon rank test for groups with various angle differences between stimulus segments for time-jittered spike trains, rounded to 2 decimal. * stands for p < 0.05.

| angle difference | Wilcoxon p |
| --- | --- |
| 90 | 0.63 |
| 68 | 0.35 |
| 45 | 0.73 |
| 23 | **0.02\*** |

Therefore, the statistical tests on time-jittered data from all three experiments showed no to very few statistically significant between-group differences. We observed no consistent pattern of significant differences depending on Gestalt structure of the stimuli across the experiments. Thus, the Gestalt structure and hence the prior probability of the stimulus can indeed be derived from the temporal structure of the data, which is captured by synchrony. When this fine temporal structure is destroyed, it is no longer possible to retrieve this information via our synchrony measure.

## Appendix

### Model parameters

| Parameter | Value | Units<br>(where applicable) |
| --- | --- | --- |
| Simulation time step | 0.005 | ms |
| Initial membrane potential | 30 | mV |
| Initial recovery variable | 30 | mV |
| Izhikevich a (time scale of recovery variable) | 0.01 |  |
| Izhikevich b (sensitivity of recovery variable) | -0.1 |  |
| Voltage afterspike reset | -65.0 | mV |
| Recovery after-spike update | 12.0 | mV |
| Excitatory reverse potential | 0.0 | mV |
| After-spike neurotransmitter presence in lateral connections | 0.02 | ms |
| After-spike neurotransmitter presence in external connections (from input to model neurons) | 0.02 | ms |
| Rising time of open receptors in lateral connections | 8.0 | ms |
| Rising time of open receptors in external connections | 8.0 | ms |
| Decay time of open receptors lateral connections | 8.0 | ms |
| Decay time of open receptors in external connections | 8.0 | ms |
| Maximum transmitter concentration in lateral connections | 1.0 |  |
| Maximum transmitter concentration in external connections | 1.0 |  |
| Voltage noise amount | 0.3 |  |

### Input parameters

| Parameter | Value | Units<br>(where applicable) |
| --- | --- | --- |
| Stimulus width | 20 | pixels |
| Stimulus height | 20 | pixels |
| Angle resolution | 22.5 | degrees |
| Input firing rate | 40.0 | Hz |
| Input noise amount | 0.4 |  |
| Input noise type | Gauss |  |
| Input strength | 2.2 |  |

### Connectivity parameters

| Parameter | Value | Units<br>(where applicable) |
| --- | --- | --- |
| Spatial distance weight | 0.5 |  |
| Orientation difference weight | 0.5 |  |
| Lateral connection weight | 2.2 |  |

### Simulation parameters

| Parameter | Value | Units<br>(where applicable) |
| --- | --- | --- |
| Simulation length | 2000 | ms |
| Downsampling | 100 | timesteps |

## Notes

### Competing Interest Statement

The authors have declared no competing interest.

